# Analyses of trait evolution and diversification reveal multiple adaptive periods in the fungal order *Hymenochaetales*

**DOI:** 10.1101/2024.06.06.597693

**Authors:** Xue-Wei Wang, Torda Varga, Qiu-Shi Li, László G. Nagy, Li-Wei Zhou

**Affiliations:** State Key Laboratory of Mycology, Institute of Microbiology, Chinese Academy of Sciences, Beijing 100101, P.R. China; University of Chinese Academy of Sciences, Beijing 100049, P.R. China; Comparative Fungal Biology, Royal Botanic Gardens, Kew, Richmond, TW9 3AB, United Kingdom; Synthetic and Systems Biology Unit, Institute of Biochemistry, Biological Research Center, HUN-REN, Temesvári krt. 62, Szeged H-6726, Hungary

**Keywords:** *Basidiomycota*, diversification rate, adaptive radiation, morphology, parasites, global warming, the expansion of angiosperms

## Abstract

Mushroom-forming fungi (*Agaricomycetes*) represent one of the most speciose and morphologically diverse life forms, which radiated into most niches on Earth and evolved diverse morphologies and life histories. The order *Hymenochaetales* comprises a species-rich group of important wood decayers and tree parasites, however, patterns of macroevolution across the order and thus the origins of key traits, such as pathogenicity are currently unknown. Here, using a novel, nearly comprehensive phylogeny of the *Hymenochaetales*, we show that its evolution has been shaped by an early adaptive period, followed by a general rate slowdown interrupted by local rapid radiations in the Cretaceous. We inferred that the ancestor of the order has undergone rapid phenotypic diversification into a range of morphologies and lifestyles, of which crust-like morphologies and ‘pileate-sessile’ forms typical of bracket-fungi became dominant among extant taxa. Net diversification rate showed significant correlations with paleoclimate, morphological and ecological traits, however, teasing apart the contributions of individual clades revealed that these were mostly driven by the Mid-late Cretaceous rapid radiation of the genus *Phylloporia*, a diverse group of plant parasites. Together, this study unraveled a complex evolutionary history of the *Hymenochaetales* and uncovered novel patterns of phenotype evolution and diversification in mushroom-forming fungi.

## INTRODUCTION

It is estimated that there are 2.2 to 3.8 million species of fungi, making them the second most speciose group of organisms after insects (Purvis & Hector 2000, Hawksworth & Lücking 2017). This species diversity is coupled with similarly varied morphologies and lifestyles. In terms of morphology, sexual fruiting bodies, which primarily support above-ground spore dispersal, stand out as some of the most spectacular and complex structures known in the fungal world. Fruiting bodies reached their highest complexity in the *Agaricomycetes* and include crust-like (resupinate) and coral-like (coralloid) morphologies, bracket fungi (pileate-sessile) or the well-known form with cap and stipe (pileate-stipitate) (Nagy et al. 2017). In addition, fungi are responsible for key ecological processes, such as the breakdown of organic matter and the growth and survival of other organisms, including plants and animals in a wide range of environments (e.g. terrestrial and aquatic habitats, Peay et al. 2016). It is conceivable that trends and patterns in the evolution of fruiting body morphologies and lifestyles can influence the rate of species birth and extinction, referred to as diversification rate, however, these are currently poorly known.

Previous studies have investigated evolutionary transitions between both fruiting body types and lifestyles by ancestral character state reconstructions in a phylogenetic framework. Resupinate morphologies and a saprotrophic lifestyle have been inferred as the ancestral condition in the *Agaricomycetes* (Hibbett & Binder 2002, Hibbett 2004, Binder et al. 2005, Hibbett 2007, Varga et al. 2019, Sánchez-García et al. 2020), from which other morphologies and lifestyles derived. Transitions to pileate-stipitate basidiomes correlated with higher diversification rates, suggesting this morphology represents a key innovation in the *Agaricomycetes*, possibly due to its increased spore-producing capacity (Varga et al. 2019, Varga et al. 2022). At the same time, mycorrhizal lifestyle may have impacted diversification rates at local phylogenetic scales (Sánchez-García et al. 2020, Sato 2024). While previous studies identified broad trends across the *Agaricomycetes*, they were based on incompletely sampled class-level phylogenies, which may not provide accurate estimates in specific lineages. It is well known that analyses of diversification rates and ancestral state reconstruction depend on sampling density, and that incomplete or, in particular, unbalanced datasets can lead to spurious inferences (Ricklefs 2007, Scholl & Wiens 2016, Title & Rabosky 2019). Therefore, accurately recovering trends in morphological and lifestyle evolution in mushroom-forming fungi requires higher sampling density and focus on specific key orders.

The *Hymenochaetales* is one of the largest orders of mushroom-forming fungi, and includes more than 1,300 described species (Wang et al. 2021, 2023, Wu et al. 2022). This fungal order is famous for accommodating some precious medicinal mushrooms (Jiang et al. 2020, Zhou et al. 2022, Cheng et al. 2023) and destructive forest pathogens (Wang et al. 2022). The species in *Hymenochaetales* show prominent diversity in both morphological and ecological traits (e.g. Korotkin et al. 2018, Zhang et al. 2018, Wang et al. 2021, 2023, Wu et al. 2022). For example, the morphology of basidiomes varies from relatively simple, resupinate to more complex, pileate-sessile (with lateral cap and no stalk), pileate-stipitate (with cap and stalk) and coralloid forms (Fig. 1A–D). The morphology of hymenophores (spore-bearing surface) shows a graded series of complexity levels, including smooth, granulated (grandinioid), toothed (odontioid to hydnoid), or those with pores (poroid) and gills (lamellate) (Fig. 1E–H). Ecologically, *Hymenochaetales* is mainly composed of saprotrophic wood-inhabiting fungi that prefer angiosperms as hosts, while some are parasites and symbionts and grow on gymnosperms or mosses (Tedersoo et al. 2007, Korotkin et al. 2018, Wu et al. 2022).

**Fig. 1.**
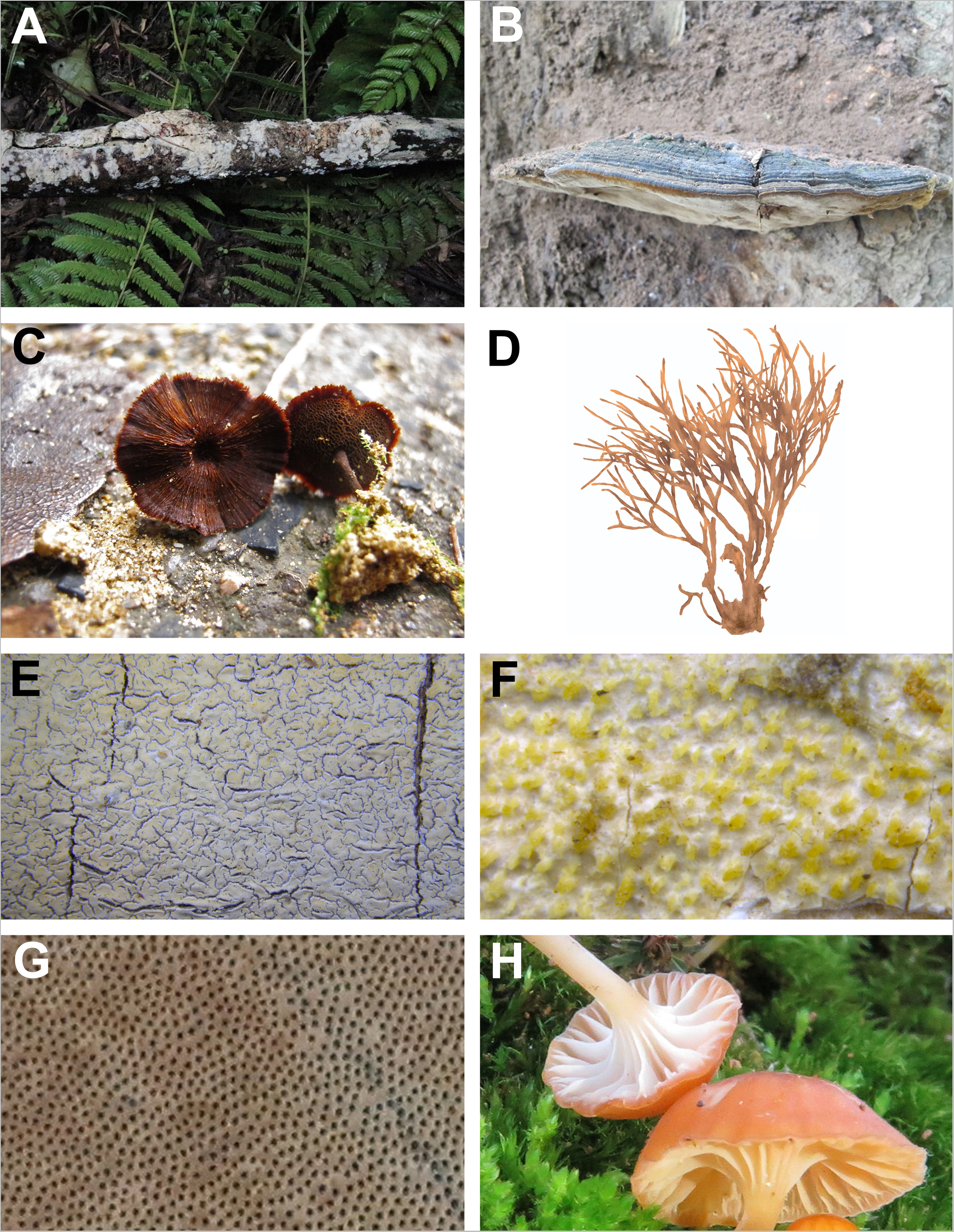
Morphotypes of basidiomes and hymenophores divided in *Hymenochaetales.* Basidiomes: resupinate (*Xylodon nesporii*, **A**), pileate-sessile (*Fulvifomes thailandicus*, **B**), pileate-stipitate (*Coltricia* sp., **C**) and coralloid (*Clavariachaete rubiginosa*, drawing from the photos in Larsson et al. 2006, **D**) forms. Hymenophores: smooth to granulated (*Schizocorticium lenis*, **E**), toothed (*Skvortzovia qilianensis*, **F**), poroid (*Sanghuangporus weigelae*, **G**) and lamellate (*Rickenella danxiashanensis*, **H**) forms.

Many previous studies made efforts to clarify the species diversity and phylogenetic relationships of *Hymenochaetales* (Zhou 2015, Zhou et al. 2018, Liu et al. 2019, Wang et al. 2020, Shen et al. 2021, Yu et al. 2021) partially due to many species in this order possessing significant commercial potentials (Wu et al. 2019) and ecological functions (Tedersoo et al. 2007, Cui et al. 2015, Wang et al. 2022). Wang et al. (2021) explored the evolution of different forms of basidiomes and hymenophores in *Hymenochaetales*; however, this study focused mainly on species in *Hyphodontia* sensu lato. In addition, it remains a major challenge to understand how the trophic modes and host preferences of *Hymenochaetales* influenced their evolutionary history. A robust phylogenetic framework is crucial for exploring the evolution of biological traits (Zhou & May 2023). Recently, Wang et al. (2023) reconsidered the taxonomy of *Hymenochaetales* accepting 14 families containing 64 genera, and another 19 genera independent from these families. Soon after, Wang & Zhou (2024) further defined the fifteenth family *Umbellaceae* with the new genus *Umbellus* as the type genus.

In this study, we reconstructed a phylogeny comprising a 99.6% of the known species in the *Hymenochaetales* and used it to analyze macroevolutionary patterns of morphological and ecological traits as well as the impact of these traits on diversification. We inferred a complex scenario involving diversification rate slowdowns, accelerations corresponding to rapid radiations, as well as adaptive periods with specific footprints of morphological and ecological trait evolution. We inferred that resupinate and pileate-sessile basidiomes, poroid hymenophores, a saprotrophic trophic mode and host preference of angiosperms became the dominant in the *Hymenochaetales*. Our results also support a rapid radiation of the genus *Phylloporia*, which might represent a period of swift adaptive species birth.

## MATERIALS AND METHODS

### Phylogeny

The internal transcribed spacer (ITS) and nuclear large ribosomal subunit (nLSU) regions are the most commonly applied phylogenetic markers for *Hymenochaetales* (Wang et al. 2023). Therefore, to sample taxa as comprehensively as possible, all species in *Hymenochaetales* with available ITS and nLSU gene sequences were selected from various references (e.g., Wang et al. 2021, 2023, Wu et al. 2022) for phylogenetic analysis (Table S1). The ITS and nLSU sequences were simultaneously used to represent one species only when they were generated from the same individual indicated by the same voucher number of collection. In addition to taxa in *Hymenochaetales*, *Fomitopsis pinicola* and *Grifola frondosa* from the *Polyporales*, and *Boletopsis leucomelaena* and *Thelephora ganbajun* from the order *Thelephorales* were selected as outgroup taxa according to Hibbett et al. (2007).

ITS and nLSU regions were separately aligned using MAFFT v.7.110 (Katoh & Standley 2013) under the “G-INS-i” option (Katoh et al. 2005). Then, the two alignments were concatenated. Misidentified sequences were filtered out from the concatenated alignment by manually inspecting preliminary phylogenetic trees inferred by maximum likelihood (ML) algorithm using raxmlGUI v.8.2.12 (Stamatakis 2014) with the inference of bootstrap (BS) replicates under the auto FC option (Pattengale et al. 2010). The best-fit evolutionary model of the concatenated alignment for ML phylogenetic analyses was estimated using jModelTest v.2.1.10 (Darriba et al. 2012, Guindon & Gascuel 2003) under the Akaike information criterion. When taxa were placed in unexpected or conflicting phylogenetic position at the genus or family level, the sequences representing these taxa were replaced by alternative sequences of the same taxa or directly removed if alternative sequences were unavailable. Eventually, a misidentification-free concatenated alignment of ITS and nLSU regions (File S1) was used to reconstruct a ML tree, with the multilocus ML tree generated in our previous study (Wang et al. 2023) as a backbone to constrain deep nodes.

Based on the resulting ML tree, an ultrametric tree with divergence times was generated via the penalized likelihood method using treePL v.1.0 (Smith & O’Meara 2012). In treePL, the value of 1000 was selected as the smoothing parameter through random-subsample-and-replicate cross-validation from values between 0.1 and 1000, while a total of 15 secondary calibration points were set on the basis of previously estimated divergence times of 14 families in *Hymenochaetales* and the origin of *Hymenochaetales* itself (Wang et al. 2023). The crown age of *Hymenochaetales* was fixed as 228 million years (Myr), while the crown age of 14 accepted families in *Hymenochaetales* were set according to their 95% highest posterior density, viz. 72-152 Myr for *Chaetoporellaceae*, 140-183 Myr for *Hymenochaetaceae*, 12-47 Myr for *Hyphodontiaceae*, 45-150 Myr for *Odonticiaceae*, 54-113 Myr for *Peniophorellaceae*, 38-107 Myr for *Repetobasidiaceae*, 39-108 Myr for *Resiniciaceae*, 15-49 Myr for *Rickenellaceae*, 120-187 Myr for *Rigidoporaceae*, 62-139 Myr for *Schizocorticiaceae*, 98-150 Myr for *Schizoporaceae*, 90-161 Myr for *Sideraceae*, 115-183 Myr for *Skvortzoviaceae* and 70-146 Myr for *Tubulicrinaceae* (Wang et al. 2023).

To avoid the possible inaccurate evolutionary reconstruction from an incompletely sampled phylogeny, a stochastic polytomy resolution method, Taxonomic Addition for Complete Trees (TACT) was used to generate an almost fully sampled phylogeny (Chang et al. 2020). TACT employs birth–death-sampling estimators across an ultrametric phylogeny to estimate branching times for unsequenced taxa with reference to their taxonomic positions, and then compatibly places the unsequenced taxa into the above-generated ultrametric tree. The known species in *Hymenochaetales* were collected according to the latest references (e.g. Wang et al. 2021, Wu et al. 2022, Wang et al. 2023) and Index Fungorum (https://www.indexfungorum.org/Names/Names.asp; accessed before June 1, 2023). The nomenclatural synonyms were considered to be the same taxa and thus only represented by their prior names. Moreover, up to June 1, 2023 the newly published taxa at the family and genus levels were also included, whereas a few species published after that time were omitted.

In the current study, except *Kurtia* and *Subulicium*, from which reliable sequence data is unavailable (Wang et al. 2023), 82 out of 84 genera in *Hymenochaetales* accommodating 99.6% of known species (1330 out of 1336 species) were included in the final time-calibrated phylogeny. In subsequent analyses of trait evolution and net diversification rate, the four species from *Polyporales* and *Thelephorales* were removed from the ultrametric tree using the R package ape v.5.7.1 (Paradis & Schliep 2019).

### Character coding of traits

Four traits, basidiome and hymenophore morphology, trophic mode and host plant were retrieved from various references (Table S1) and subjected to evolutionary analyses.

The basidiomes were divided into four different morphotypes: 1) resupinate (721 taxa), 2) pileate-sessile (463 taxa), 3) pileate-stipitate (142 taxa), and 4) coralloid (four taxa) (Table S1). The resupinate form was defined as taxa that lay flat on the substrate, such as crust-like, effused-reflexed, ceraceous and membranous forms. Pileate-sessile forms accommodated taxa bearing a cap, but lacking a stipe or only having a rudimentary or reduced stipe. Forms that have a well-defined cap and a stipe were considered as pileate-stipitate. Club-shaped and branched forms were considered to be coralloid.

The hymenophores were coded as 1) poroid (835 taxa), 2) smooth to granulated (353 taxa), 3) toothed (126 taxa), or 4) lamellate (16 taxa) (Table S1). Poroid type indicated species with porous hymenophoral configuration. Smooth to granulated type was used to define species with smooth or tuberculate hymenophoral surfaces. Species with clearly toothed surface were designated as toothed type. Lamellate type was defined as the presence of thin and pleated hymenophoral configuration.

Three different trophic modes were distinguished: 1) saprotrophs (1113 taxa), 2) parasites (151 taxa), and 3) symbionts (61 taxa) (Table S1). When no clear record about trophic modes is available, species found on dead trees, decayed or burnt wood, burnt soil (humus), litter and dung were treated as saprotrophs, and those growing on living parts of trees were treated as parasites. Previous research clarified that in *Hymenochaetales* most of taxa growing on ground were symbionts (Tedersoo et al. 2007, Korotkin et al. 2018), so accordingly we treated species on ground as symbionts (Tedersoo et al. 2007, Korotkin et al. 2018). In addition, species (five taxa) lacking information for being assigned to any trophic mode were treated as missing data.

Hosts were generally separated as: 1) angiosperms (1098 taxa), 2) gymnosperms (163 taxa), 3) both angiosperms and gymnosperms (47 taxa), and 4) mosses (22 taxa) (Table S1). For species growing on the ground, their hosts were designated as either angiosperm or gymnosperm on the basis of the spatially close dominant plants.

### Ancestral trait reconstruction and trait transition

The evolutionary histories of traits were inferred using the R package CORHMM v.2.8 (Beaulieu et al. 2013), which can handle polymorphic tips and missing values. Firstly, the ancestral trait was reconstructed jointly using the ML estimation with an “all-rates different” model of the function ‘corHMM’, which could give one best global estimate for each ancestral node. By assuming that a transition between two adjacent nodes could happen once at any time along the branch connecting them, the average number of transitions through time and across branches was calculated using a 12 Myr time window with the step size of 6 Myr. Then, the number of transitions in internal nodes exclusively, as well as both internal and terminal nodes were calculated separately with the help of a custom R script (available at https://github.com/vtorda/TraitEvolution and from the authors on request). Furthermore, to explore the dynamics of trait transition events across branches and to estimate the posterior probability distribution for the most recent common ancestor (MRCA) of each node, the evolutionary histories of traits were randomly mapped on the time-calibrated ultrametric tree 100 times using the function ‘makeSimmap’ and the set of stochastic maps were summarized. The tree plot was created using the function ‘ggtree’ of the R package ggtree v.3.6.2 (Yu 2020). The posterior probability distribution for the MRCA of each node was plotted as a pie chart using the function ‘nodelabels’ of the R package ape v.5.7.1 (Paradis & Schliep 2019).

### Trait-independent diversification analysis

Bayesian Analysis of Macroevolutionary Mixtures (BAMM, Rabosky 2014) was used to estimate rate heterogeneity across lineages and detect shifts in diversification rates independent from character states. The approach assumes that different diversification regimes occurred across the branches of a phylogenetic tree, and accounts for rate variation through time and among lineages. It provides estimates of the number of diversification-rate shifts across the branches of a tree and estimates diversification-rate parameters. The analysis was run for 100 million Markov chain Monte Carlo (MCMC) generations using four independent chains and sampling parameters every 10,000 generations. Ten thousand of the posterior samples were stored, with 2,500 discarded as burn-in and leaving 7,500 samples for subsequent analysis. The convergence of MCMC was checked with an effective sample size above 200 by the R package CODA v.0.19.4 (Plummer et al. 2006) after discarding the first 25% of posterior samples as burn-in. The R package BAMMtools v.2.1.10 (Rabosky et al. 2014) was used to estimate speciation and extinction priors with the “setBAMMpriors” function, and also to evaluate the outputs. In BAMM, the diversification analysis was run three times with each generating almost identical results, and thus only the results of the first run are shown.

### Correlation between paleotemperature and the net diversification rate

The correlation between the trend of paleotemperature and the net diversification rate curve of *Hymenochaetales* was carried out by a custom R script of Davis et al. (2016) with adapted revisions for our data. The paleotemperature reflected through the smoothed δ^18^O data was taken from Davis et al. (2016), while the net diversification rate of *Hymenochaetales* was generated from BAMM. Briefly, the Pearson’s cross-correlation analysis (PCCA) was performed for a linear correlation analysis. Given the normal distribution obtained, a Student’s *t*-test was performed to assess the significance of difference of the mean from zero.

### Trait-dependent diversification analysis

Normally, the trait-dependent diversification is analyzed by the Binary-State Speciation and Extinction (BiSSE) method or Multi-State Speciation and Extinction (MuSSE) method (Maddison et al. 2007). However, due to both phylogenetic pseudoreplication and model inadequacy, these methods are prone to incorrect inference of state-dependent diversification (Maddison & FitzJohn 2015, Rabosky & Goldberg 2015). So, following the method of Chazot et al. (2021) and Hackel et al. (2022), the trait-dependent diversification analysis was conducted by randomly pairing the trait-independent diversification rate estimated by BAMM with the trait evolution stochastic maps inferred by CORHMM. In addition, a sliding time window with a range of 12 Myr and the step size of 6 Myr was used to estimate the diversification rates through time. The diversification rate per trait were then calculated for both overall time and each of 19 equal-sized time windows. Within each time window, if a lineage represented a character of the four traits, the rates estimated for this lineage will contribute to the average rate of the character. The average was computed by estimating the number of transition events of one trait (rate multiplied by branch length) divided by the sum of branch lengths involved in this trait during the same time interval. The above trait-dependent analysis was conducted using a custom R script of Hackel et al. (2022) with adapted revisions for our data.

## RESULTS

### Phylogeny

The concatenated alignment of ITS and nLSU regions accommodates 815 sequence-available species in *Hymenochaetales*, besides four species from *Polyporales* and *Thelephorales* (File S1). The resulting ML phylogenetic tree recovered all 15 accepted families and 17 incertae sedis genera in *Hymenochaetales* as monophyletic (Fig. S1) that is consistent with the latest multilocus and genome-scale phylogenetic treatments (Wang & Zhou 2024, Zhao et al. 2023), and most of the nodes received high bootstrap supports (>=70%). The divergence times of each family in the time-calibrated tree generated by treePL are similar to that of Wang et al. (2023). In general, *Hymenochaetales* originated in Triassic, which is a bit older than that inferred by Varga et al. (2019) and Zhao et al. (2023). Of the 15 families, 13 emerged during the Jurassic and Cretaceous, while *Hyphodontiaceae* and *Rickenellaceae* evolved in the early Cenozoic. Then, additional 515 unsequenced species in *Hymenochaetales* were incorporated into the time-calibrated tree using TACT. Eventually, an almost fully sampled tree with a 99.6% sampling proportion of all known species in *Hymenochaetales* was generated (Fig. S2).

### Evolutionary instability of morphological and ecological traits

To understand adaptive evolution in the *Hymenochaetales*, two morphological (basidiome and hymenophore morphology) and two ecological (trophic mode and host) traits were analyzed. Ancestral state reconstructions suggest that the MRCA of *Hymenochaetales* most likely to possess crust-like, resupinate basidiomes (posterior probability of 0.89) and a simple, smooth to toothed hymenophore configuration (see Fig. 1E–F, the smooth to granulated form had a posterior probability of 0.53, while the toothed form had a posterior probability of 0.47), and grew on seed plants (posterior probability for angiosperms & gymnosperms of 0.91) as a saprotroph (posterior probability of 0.99) (Figs. S3–S6). The morphological reconstructions are consistent with previous studies (Hibbett & Binder 2002, Hibbett 2004, Varga et al. 2019), which inferred that most of the early evolution of the *Agaricomycetes* was dominated by resupinate forms. We note that angiosperms & gymnosperms as the inferred ancestral substrate is likely a consequence of the ubiquity of generalist extant taxa, as, according to literature data, angiosperms have not yet evolved when the MRCA of *Hymenochaetales* existed.

We analyzed the temporal distribution of character state transitions in the four traits to understand the dynamics of phenotypic evolution and potential adaptive periods during the history of the *Hymenochaetales*. Because the TACT algorithm places unsequenced species based on taxonomy, which can introduce spurious transitions, we removed all tips from the analyses. When only internal nodes are considered, trophic mode is the most evolutionarily stable trait, for which 37 state transitions were inferred. The order is dominated by saprotrophs and the ancestral lifestyle of the order was inferred as saprotrophy, which evolved into symbionts and parasites on a few occasions (Figs. 2C & S5).

**Fig. 2.**
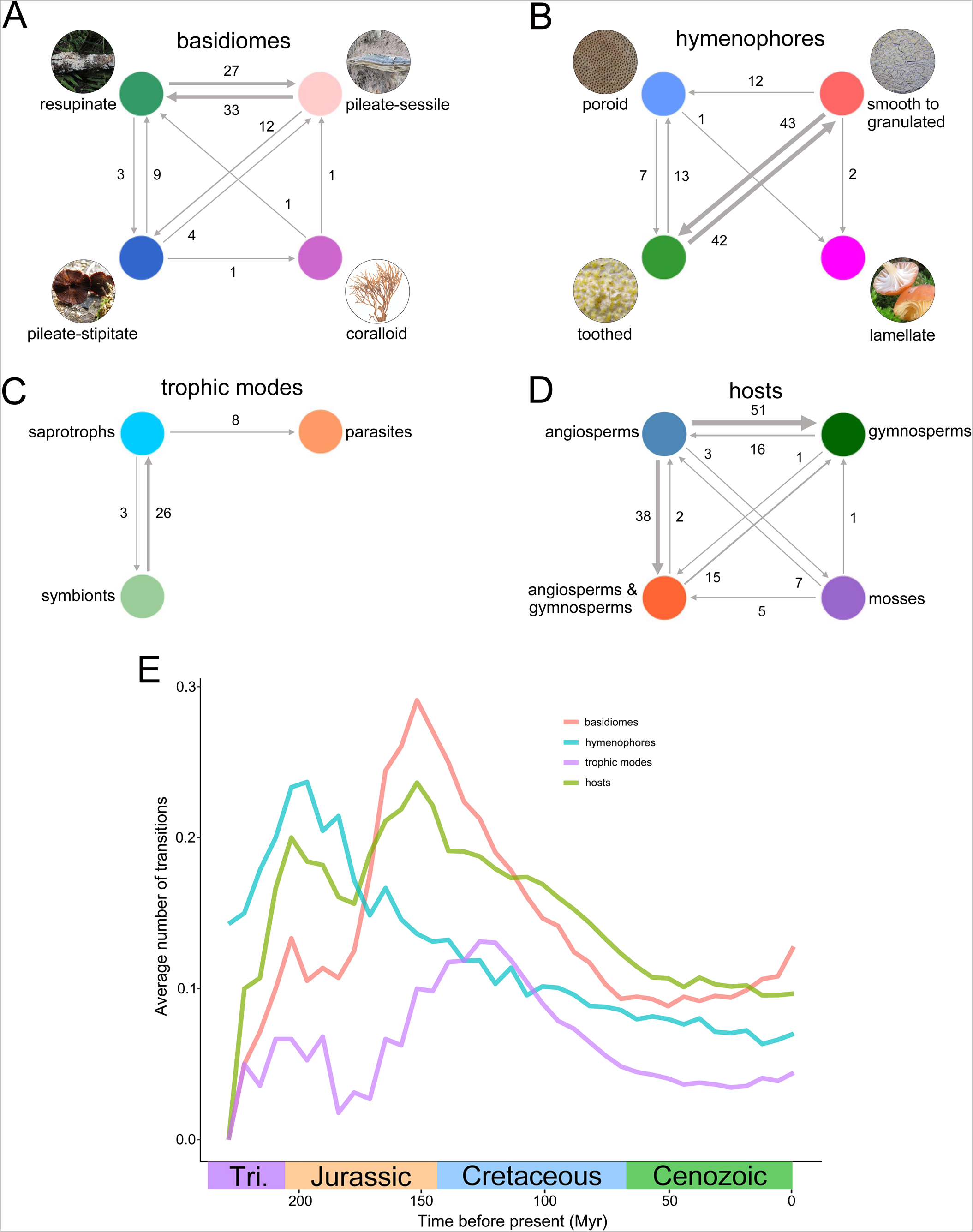
The number of transition events of internal nodes between different forms and the transition events frequency through time of each trait. (**A**) The number of transition events of basidiomes. (**B**) The number of transition events of hymenophores. (**C**) The number of transition events of trophic modes. (**D**) The number of transition events of hosts. (**E**) The transition numbers through time.

On the other hand, host association showed the largest number, a total of 139 transitions (Fig. 2D), which makes this the most labile trait examined in this study. This is followed by hymenophore morphology (110 transitions) and basidiome type (91 transitions) (Fig. 2A & B). Within a given trait, the largest number of state transitions were inferred between smooth to granulated and toothed hymenophores (85 transitions), between angiosperm and gymnosperm hosts (67 transitions) and between resupinate and pileate-sessile basidiomes (60 transitions) (Fig. 2). The transition events inferred from all nodes and only tips showed similar patterns, indicating the robustness of our results to excluding tips from the data (Fig. S7). Taken together, these data indicate that the examined traits showed a large number of transitions during the evolution of the *Hymenochaetales*, suggesting an evolutionary instability of morphologies and lifestyles in the order.

We were next interested in whether state transitions are more common at certain parts of the tree or in specific time windows, which would indicate periods of quick adaptive change. The analysis shows the frequency of state transitions across evolutionary time, in sliding windows of 12 Myr (Fig. 2E). All four traits show a peak in the early evolution of the order, which corresponds to basal branches of the tree. Basidiome types and host plants showed the highest number of transitions in the late Jurassic (26 and 21 transitions among 96 lineages, respectively, between 150 and 160 Myr ago), hymenophore in the early Jurassic (seven transitions among 30 lineages between 190 and 200 Myr ago), whereas lifestyle in the early Cretaceous (21 transitions among 156 lineages between 125 and 135 Myr ago). The most evident peak was observed in basidiome morphologies. The frequency of transitions decreased after these peaks and overall show a deceleration towards the present. This suggests that most of the state transitions concentrate to the early evolution of the order and indicates a lability of character states in this period.

These patterns are also empirically evident if we compare basal and more derived clades in the *Hymenochaetales*. A high diversity of morphologies can be found in early-diverging clades of the *Hymenochaetales*. Of the four basidiome types, coralloid and pileate-stipitate occur only in the basal clade comprising *Rickenellaceae* and associated genera, which includes typical pileate-stipitate fungi with gills (e.g. *Rickenella* and *Contumyces*), pileate-stipitate to fan-shaped with smooth hymenophores (e.g. *Cotylidia*), cupulate (e.g. *Muscinupta*), coralloid (e.g. *Alloclavaria* and *Bryopistillaria*) but also resupinate forms (e.g. *Ginnsia*, *Globulicium* and *Lyoathelia*) (see Fig. 1). The basal clade comprising *Rickenellaceae* and associated genera also harbor the highest diversity of lifestyles, including moss-associated, soil and wood saprotrophs. In comparison, basidiome morphology, hymenophoral configurations and lifestyle show higher homogeneity in other families, such as *Hymenochaetaceae*, *Hyphodontiaceae*, *Rigidoporaceae* and *Schizoporaceae*. Of these families, *Hymenochaetaceae* is particularly notable, given that in terms of species diversity this is the most diverse in the order.

### Diversification rate slowdown and rapid radiations in the Hymenochaetales

Diversification rates were inferred using the time-calibrated phylogeny in BAMM with rate shifts allowed. This revealed global diversification patterns across the *Hymenochaetales* in the last ∼200 Myr. The per-branch net diversification rate was estimated to be between 0.019 to 0.3 per Myr (Fig. 3A) and fluctuated considerably during the history of the order (Fig. 3B). The ancestor of the *Hymenochaetales* was inferred to have a high diversification rate, followed by a slowdown in the Triassic and Jurassic. Across the whole order, we inferred a sudden peak in diversification rates in the Cretaceous, ∼100 Myr ago (Fig. 3B). We identified six significant rate shifts occurring at different times and within six distinct clades (Fig. 3A). These diversification rate shifts were robust, with the top nine shift configurations containing almost the same rate shifts, but at different marginal shift probabilities (Fig. S8). Rate shift 1 occurred in late Triassic and was associated with a slowdown of net diversification rate soon after the origin of the order, consistent with the overall rate tendencies (Fig. 3B). Such slowdowns are often identified when rapidly diversifying species fill available niches during adaptive radiations, suggesting that in the early evolution of the *Hymenochaetales* diversification may have been limited by niche availability.

**Fig. 3.**
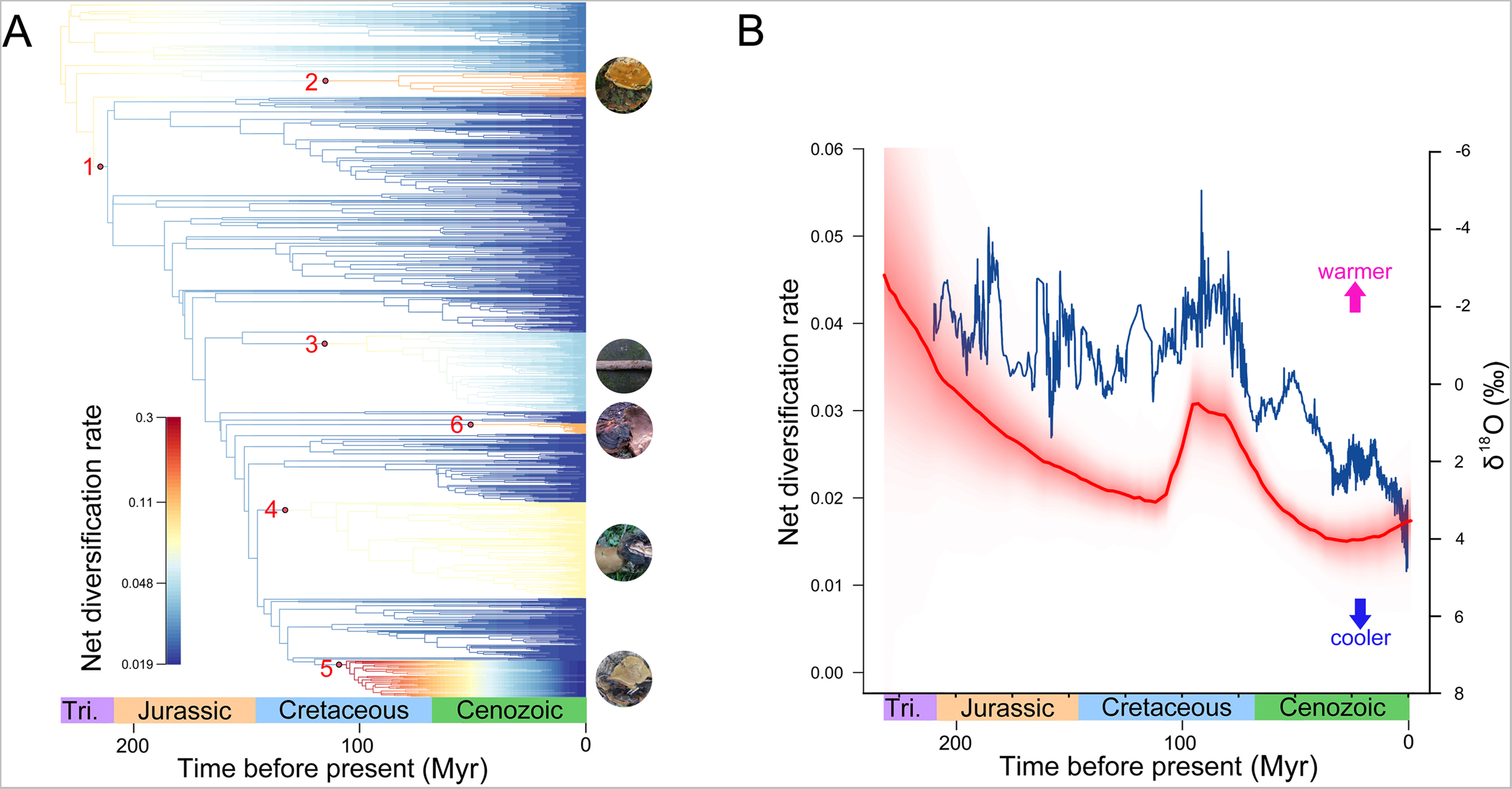
Trait-independent net diversification rate of *Hymenochaetales*. (**A**) Per-branch net diversification rate averaged across posterior samples. The rate shift 1 indicates the lineage being composed of *Chaetoporellaceae*, *Hymenochaetaceae*, *Hyphodontiaceae*, *Repetobasidiaceae*, *Schizoporaceae*, *Tubulicrinaceae*, *Hastodontia* and *Sphaerobasidium*, 2 *Rigidoporus*, 3 *Hymenochaete*, 4 the lineage consisting of *Clavariachaete*, *Inonotus*, *Meganotus*, *Ochrosporellus*, *Pachynotus*, *Perenninotus*, *Rigidonotus*, *Sanghuangporus*, *Sclerotus* and *Tropicoporus*, 5 *Phylloporia*, and 6 *Porodaedalea*. The schematic basidiome morphologies of the lineages corresponding to the shifts 2-6 are shown at the right of the tree. Root age was fixed as the previous estimate of 228 million years for the crown age of *Hymenochaetales* (Wang et al. 2023). (**B**) The δ^18^O curve (blue line) superimposed on the net diversification rate (red line) through time. Shaded red areas are 95% quantile ranges.

Rate shifts 2-5 occurred in the Cretaceous and shift 6 occurred in Cenozoic, and all were associated with an increase of net diversification rate (Fig. 3A). Rate shifts 2-5 were inferred between 150 and 100 Myr ago. These were associated with the genera *Rigidoporus* (shift 2), *Hymenochaete* (shift 3), *Clavariacheate* to *Tropicoporus* (shift 4), *Phylloporia* (shift 5) and *Porodaedalea* (Shift 6). These overlap with those reported by Varga et al. (2019), who detected five rate shifts in the *Hymenochaetales*, based on a different dataset, underscoring the dynamic diversification rates in the order. The most abrupt diversification rate increase, around 4-fold, corresponded to the genus *Phylloporia*, ca. 100 Myr ago (Figs. 3A & S9). We found that *Phylloporia* is probably responsible for the Cretaceous diversification peak (100 Myr), as removing the *Phylloporia* clade led to the disappearance of this peak (Fig. S9G). This reflects that the rapid diversification of *Phylloporia* clade was probably the main driver of the sudden increase in diversification rate of entire *Hymenochaetales* around 100 Myr.

### Diversification rates of Hymenochaetales correlate with paleoclimate

Overlaying the diversification rate patterns of *Hymenochaetales* with the global paleotemperature identified from oxygen isotope (δ^18^O) records revealed an apparent correlation between these two parameters (Figs. 3B & S10). The gradual decrease of net diversification rate since the origin of the order in the late Triassic, correlates with decreasing temperatures in this geological time period. This trend was broken by a global warming event in the mid-late Cretaceous, when a coincident abrupt increase in net diversification rate was inferred (∼100 Myr ago, Fig. 3B).

The correlations of all realizations of the diversification curve in the BAMM analysis (7,500 in total after burn-in the first 25% samples) against the global δ^18^O curve was statistically tested. For each realization, a correlation coefficient was calculated ranging from –1 (diversification rate increases along with cooler temperatures) to 1 (diversification rate increases along with warmer temperature). The temperature has no correlation with diversification rate if the coefficient is zero. When the entire *Hymenochaetales* being considered, the resulting correlation coefficient (ρPCCA(s) = 0.659±0.003, *p* < 2e–16, Fig. S10A) significantly implies that the diversification rate of *Hymenochaetales* positively correlates with temperature. Regarding solely the *Phylloporia* clade, the significant correlation also can be found between the increase of diversification rate and warmer temperature (ρPCCA(s) = 0.875±0.000, *p* < 2e–16, Fig. S10B). At the same time, when the whole *Hymenochaetales* is considered without *Phylloporia* clade, the correlation was still significant (ρPCCA(s) = 0.269±0.006, *p* < 2e–16, Fig. S10C), albeit weaker.

These analyses suggest a clear correlation between net diversification rate and paleotemperature in the *Hymenochaetales*. Although the peak in diversification rate ∼100 Myr ago, which we found to be related to the radiation of the genus *Phylloporia*, correlates most evidently with paleotemperature, the correlation exists even beyond this diversification peak. Based on these patterns, we speculate that diversification of the *Hymenochaetales* and, possibly in particular that of *Phylloporia* was at least partially driven by temperature. This effect might have been direct or indirect, clarifying this requires more research.

### Morphological and ecological traits impact diversification rate

We also examined the possibility that diversification rates are influenced by innovations in morphological and ecological traits. These analyses suggested character states correlate significantly with net diversification rates in the *Hymenochaetales* (Fig. 4). The pileate-sessile and resupinate fruiting body morphologies had almost identical, high net diversification rates, followed by the pileate-stipitate form, while coralloid forms were associated with the lowest net diversification rates (ANOVA *p*-value < 0.001, Fig. 4A). This is consistent with the observation that resupinate and pileate-sessile species are most widespread in the order, while coralloid and pileate-stipitate species are limited to certain small genera, such as *Alloclavaria* and *Rickenella*, respectively.

**Fig. 4.**
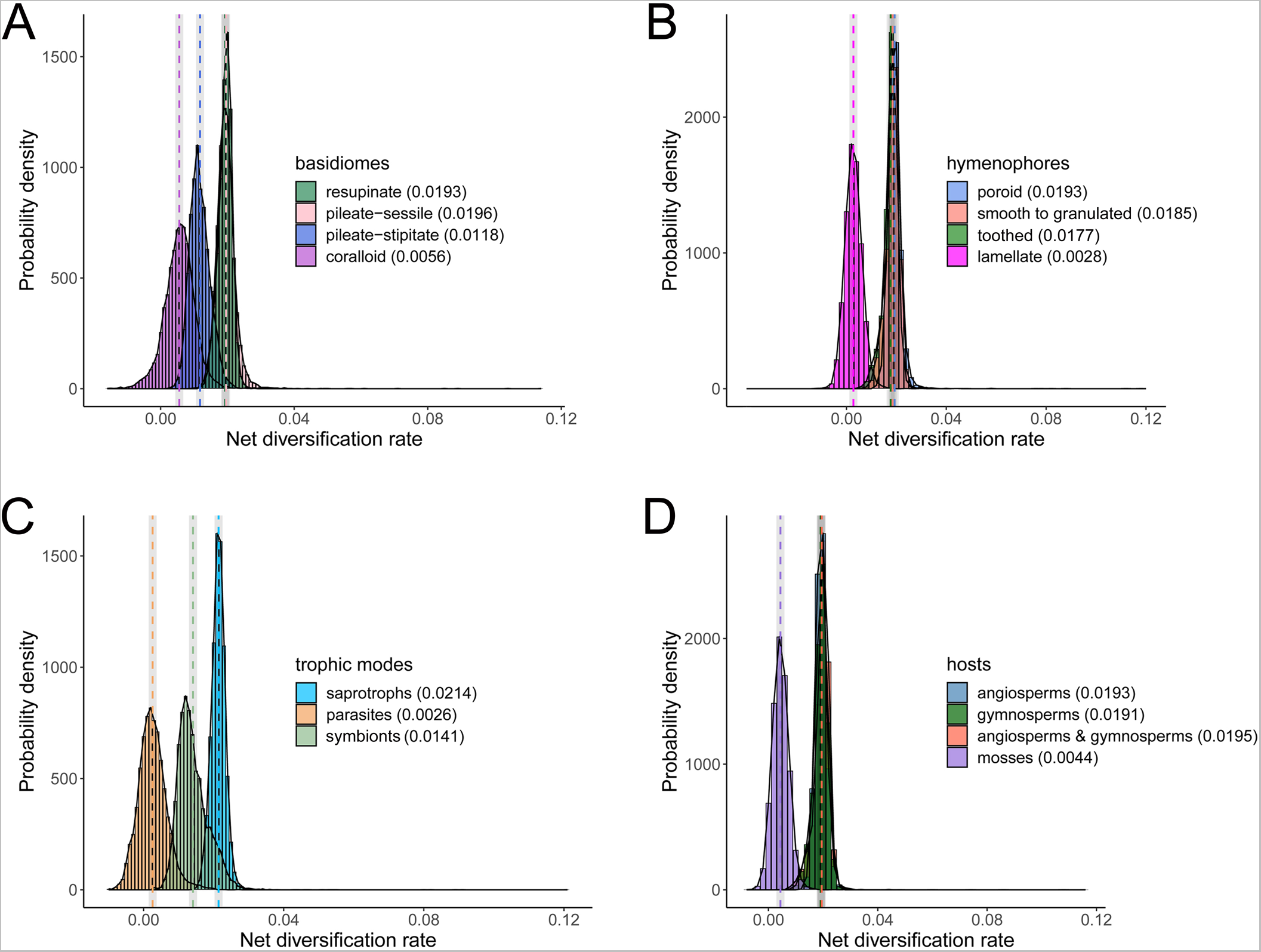
Posterior probability distribution of net diversification rates in the four traits, viz. basidiomes (**A**), hymenophores (**B**), trophic modes (**C**) and hosts (**D**). Histograms are coloured by the forms of each traits and the corresponding dashed lines represent the mean values.

Regarding hymenophores, the poroid, smooth to granulated and toothed forms had similar net diversification rates, which are higher than that inferred for the lamellate form (ANOVA *p*-value < 0.001, Fig. 4B). Net diversification rates of the saprotrophs were higher than those of the symbionts and parasites (ANOVA *p*-value < 0.001, Fig. 4C). The growth exclusively on angiosperms, on angiosperms & gymnosperms, and exclusively on gymnosperms had similar net diversification rates, while moss-associated lineages had the lowest net diversification rates (ANOVA *p*-value < 0.001, Fig. 4D).

Throughout the evolutionary history of *Hymenochaetales*, the various characters in the four traits contributed to the net diversification rate with different trends (Fig. 5). The species with ancestral characters of the four traits, viz. resupinate basidiomes, smooth to granulated hymenophores, saprotrophic trophic mode and growth on angiosperms & gymnosperms had a gradual decrease of net diversification rates since the origin of *Hymenochaetales* (Fig. 5). In contrast, the diversification rates of species with pileate-sessile basidiomes, poroid hymenophores, parasitic trophic mode and growth on angiosperms showed an increase around 100 Myr ago (Fig. 5). This trend is consistent with the trait-independent global net diversification rates (Fig. 3B). Furthermore, among all characters of the four traits, the parasitic trophic mode contributed the highest increase of net diversification rate around 100 Myr ago (Fig. 5). These increases are consistent with the radiation of the genus *Phylloporia*, which includes pileate-sessile species with a poroid hymenophore that parasitize angiosperms (Wu et al. 2022).

**Fig. 5.**
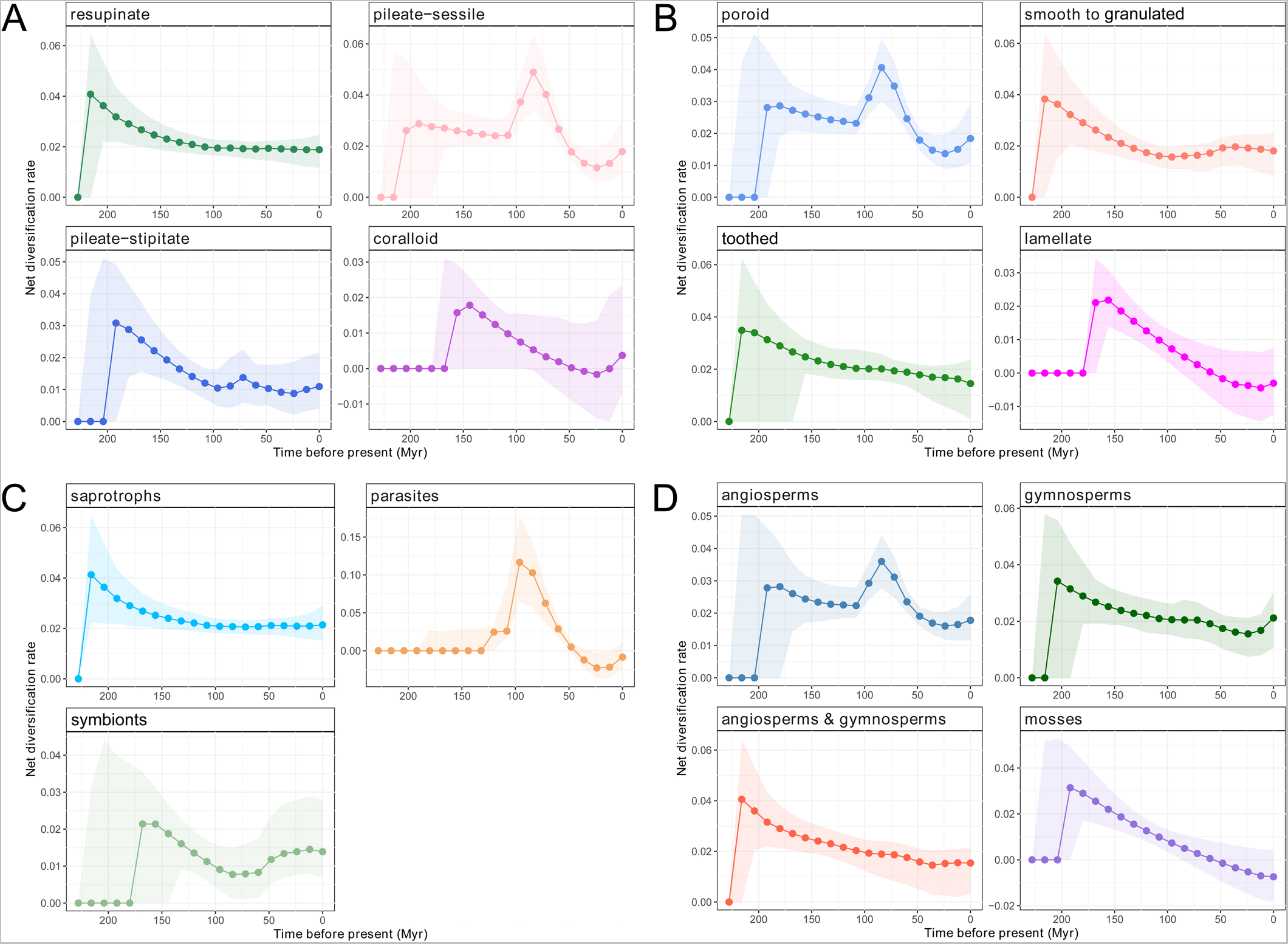
The net diversification rate of each form in the four traits, viz. basidiomes (**A**), hymenophores (**B**), trophic modes (**C**) and hosts (**D**) of *Hymenochaetales* through time.

## DISCUSSION

In this study, we examined the evolution of the *Hymenochaetales*, one of the most economically and ecologically important macrofungal orders, in the context of trait evolution and species diversification. Our analyses are based on an almost comprehensively sampled and well-supported phylogeny of 815 species supplemented with another 515 unsequenced species, which also provides a general phylogenetic framework for further research on the order. Maximizing species coverage necessitated limiting the data matrix to the ITS and nLSU regions, two widely used phylogenetic markers in fungi, which we combined with multilocus backbone tree to help the resolving ancient nodes (Wang et al. 2023). The phylogeny we inferred (Fig. S1) is consistent with the recently updated framework of *Hymenochaetales* based on multiloci (Wang et al. 2023) and phylogenomic analyses (Zhao et al. 2023, Varga et al. 2019). This phylogeny allowed us to infer broad trends in morphological and lifestyle evolution, as well as patterns of diversification in the *Hymenochaetales*.

Broad trends of trait evolution across the *Agaricomycetes* or *Basidiomycota* have been a subject of intense research (Hibbett & Donoghue 2001, Hibbett & Binder 2002, Hibbett 2004, 2007) with the recent publication of large-scale studies for resolving global patterns (Varga et al. 2019, Sánchez-García et al. 2020, Virágh et al. 2022). While these studies recovered general evolutionary trends, they did not test whether these trends generally exist in all clades within the *Agaricomycetes* and what lineage-specific processes have contributed to the diversity of the class. In this study, we identify the *Hymenochaetales* as an exception, which follows different trends from that of the *Agaricomycetes*, both in trait evolution and net diversification rates. We reconstructed the ancestor of the *Hymenochaetales* as a species that possessed resupinate basidiomes, simple, smooth to toothed hymenophoral configuration and was a saprotroph of seed plants. Our analyses further uncovered a large dynamics in character state transitions, indicating that several character states have been labile during the history of the *Hymenochaetales*. These data indicated that transitions between character states were particularly frequent in the early evolutionary history of the *Hymenochaetales*: state change frequencies of all four traits peaked early, from the Triassic to the Jurassic periods. This might be a hallmark of a period of rapid phenotypic innovation, where novel morphologies and ecologies evolved quickly after the emergence of the *Hymenochaetales* clade. This period was, based on our inferences, followed by a deceleration of morphological innovation and diversification rate, which is evident from the downwards slope of the state change frequency curves on Fig. 2E. Rapid phenotypic divergence, followed by a deceleration, is a hallmark of adaptive radiations (Glor 2010, Moen & Morlon 2014). We speculate that these evolutionary patterns established a large diversity of morphologies and ecologies in the early evolution of the *Hymenochaetales* clade. In the subsequent time periods evolution may have sorted this diversity out so that only a few phenotypes became really successful (e.g. resupinate or pileate-sessile taxa and saprotrophy), whereas others became less frequent (e.g. pileate-stipitate form).

In addition to trait evolution, patterns of the emergence and extinction of species, collectively known as diversification, have recently become a subject of intense research in mushroom-forming fungi (e.g. Wilson et al. 2011, Nagy et al. 2012, Sato et al. 2017, Sato & Toju 2019, Varga et al. 2019, Sanchez-Garcia et al. 2020) and hold the promise of identifying key innovations and the drivers of morphological and ecological diversity and species richness. The rate of diversification can vary through time and across clades, which can be indicative of rapid radiations, interactions with the environment and differences in the evolvability or evolutionary success of clades (Ricklefs 2007, Scholl & Wiens 2016). In this study we focused on diversification rate in the *Hymenochaetales* and found distinct fluctuations throughout its evolution. At the earliest nodes of the order, we inferred high diversification rates, followed by a strongly supported shift to lower rates. This rate slowdown, in addition to patterns of phenotype evolution (see above), provides further support to the hypothesis that the *Hymenochaetales* has undergone an early adaptive radiation, which has filled available niches and thus was followed by a deceleration. We identified another five significant rate shifts, which uniformly represent rate acceleration. These partially correspond to the rate shift inferred by Varga et al. (2019), however, provide more precision as to their placement, due to the nearly comprehensively sampled phylogeny we used. The most remarkable rate shift of these is the radiation of the genus *Phylloporia*, a diverse genus of pileate-sessile plant parasites that our data suggest have undergone a rapid radiation ∼100 Myr ago. This might represent a rapid radiation into niches opened by either internal or external factors, such as climate or the radiation of angiosperm host trees. Although this needs more research, if verified it would represent the second documented explosive radiation event in the fungi, after the genus *Coprinellus* (Nagy et al. 2012).

It was postulated that the pileate-stipitate basidiome morphology is a very successful adaptation in the *Agaricomycetes*, that evolved several times convergently and is found in the majority of extant species of the class (Varga et al. 2019, 2022). In the *Hymenochaetales* our results do not suggest that the pileate-stipitate morphology became a particularly successful state. Pileate-stipitate species (including classic bracket fungi), although evolved several times in the *Hymenochaetales*, were associated with low net diversification rates and did not become the dominant form. Among extant species, it is found in ∼10% of the species, including, for example, *Coltricia* and *Rickenella*. On the other hand, resupinate (54.2% 721 of 1330) and pileate-sessile (34.8% 463 of 1330) morphologies dominate the extant diversity of the order and were inferred to be associated with slightly higher diversification rates. The scarcity of pileate-stipitate species may be because it evolved only in small clades, relatively recently, or because the saprotrophic, wood-associated lifestyle of *Hymenochaetales* matches better with resupinate and pileate-sessile morphologies.

Across the character states of the four traits, we found that the pileate-sessile basidiome morphology, poroid hymenophoral configuration, parasitic lifestyle and angiosperm hosts contributed most clearly to the dynamics of diversification rates. Most of the effect was inferred around 100 Myr ago, coinciding with rate shift 5, which represents the radiation of *Phylloporia*. This genus contains species with the exact traits mentioned above, suggesting that these analyses, in fact, largely picked up the rapid radiation of *Phylloporia*. With regard to trophic modes, in *Phylloporia* parasitism might have contributed to the increase of net diversification rate around 100 Myr ago, although it is hard to separate its effect from that of other traits. Nevertheless, this is consistent with the suggestion that trophic mode impacts diversification rates at local phylogenetic scales (Sánchez-García et al. 2020, Sato 2024).

Our analyses uncovered a significant correlation between paleoclimate and diversification rate, including an increase of net diversification rate approximate 100 Myr ago in the Mid-late Cretaceous corresponds to the global warming (Ziegler et al. 2003). Although a significant portion of the signal likely came from the rapid radiation of *Phylloporia*, the correlation remained significant even after removing *Phylloporia* from the analyses. This suggests that global temperature trends, and certainly other biotic factors, may contribute to the diversification of *Hymenochaetales*. Among others, the expansion of angiosperms in the Cretaceous (Niklas et al. 1983) may be an important contributor, as *Hymenochaetales* are decayers or parasites of woody plants. This biotic factor has also been applied for explaining the diversification of *Agaricomycetes* (Krah et al. 2018, Varga et al. 2019) and other life forms (McKenna et al. 2009) feeding on plants. Therefore, it is possible that the habitat provided by angiosperms provided niches that drove the diversification of *Hymenochaetales*, among other factors.

## CONCLUSIONS

Taken together, this study provided evidence for adaptive periods in the evolution of the *Hymenochaetales*, that involved both phenotypic evolution and rates of species diversification. Our inferences suggest that the emergence of the order was followed by rapid evolution of morphologies, a deceleration of diversification rates in the Triassic/Jurassic periods, followed by five diversification rate shifts in smaller clades, of which the genus *Phylloporia* might represent a particularly rapid radiation. The data suggest that the *Hymenochaetales* had a dynamic evolutionary history that was shaped by interactions between the environment (e.g. paleoclimate), morphological and ecological innovations, as well as potential other factors not analyzed in this study. Although the *Hymenochaetales* mirrors several phenomena described globally for the *Agaricomycetes*, we also uncovered several unique patterns, indicating that analyzing trait evolution and diversification at finer phylogenetic scales is necessary to uncover the full complexity of the evolution of mushroom-forming fungi.

## ACKNOWLEDGEMENTS

The research was financed by the National Key Research and Development Program of China (No. 2022YFC2601200), and the National Natural Science Foundation of China (Project Nos. 31970012, 32111530245 & 31570014). LGN appreciates funding from the Hungarian Academy of Sciences (LP2019-13/2019) and the National Research Development and Innovation Office (Grant No. OTKA 142188).

## Supplementary materials

**Fig. S1.**
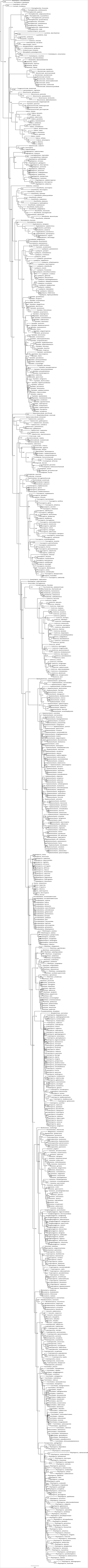
The phylogenetic tree of *Hymenochaetales* generated by maximum likelihood algorithm. Bootstrap values are indicated at the nodes.

**Fig. S2.**
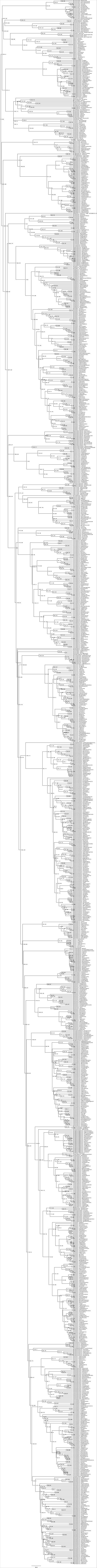
The time-calibrated ultrametric supertree with a 99.6% sample proportion of all known species in *Hymenochaetales*. The mean divergence times of clades (crown ages) were labeled at the nodes.

**Fig. S3.**
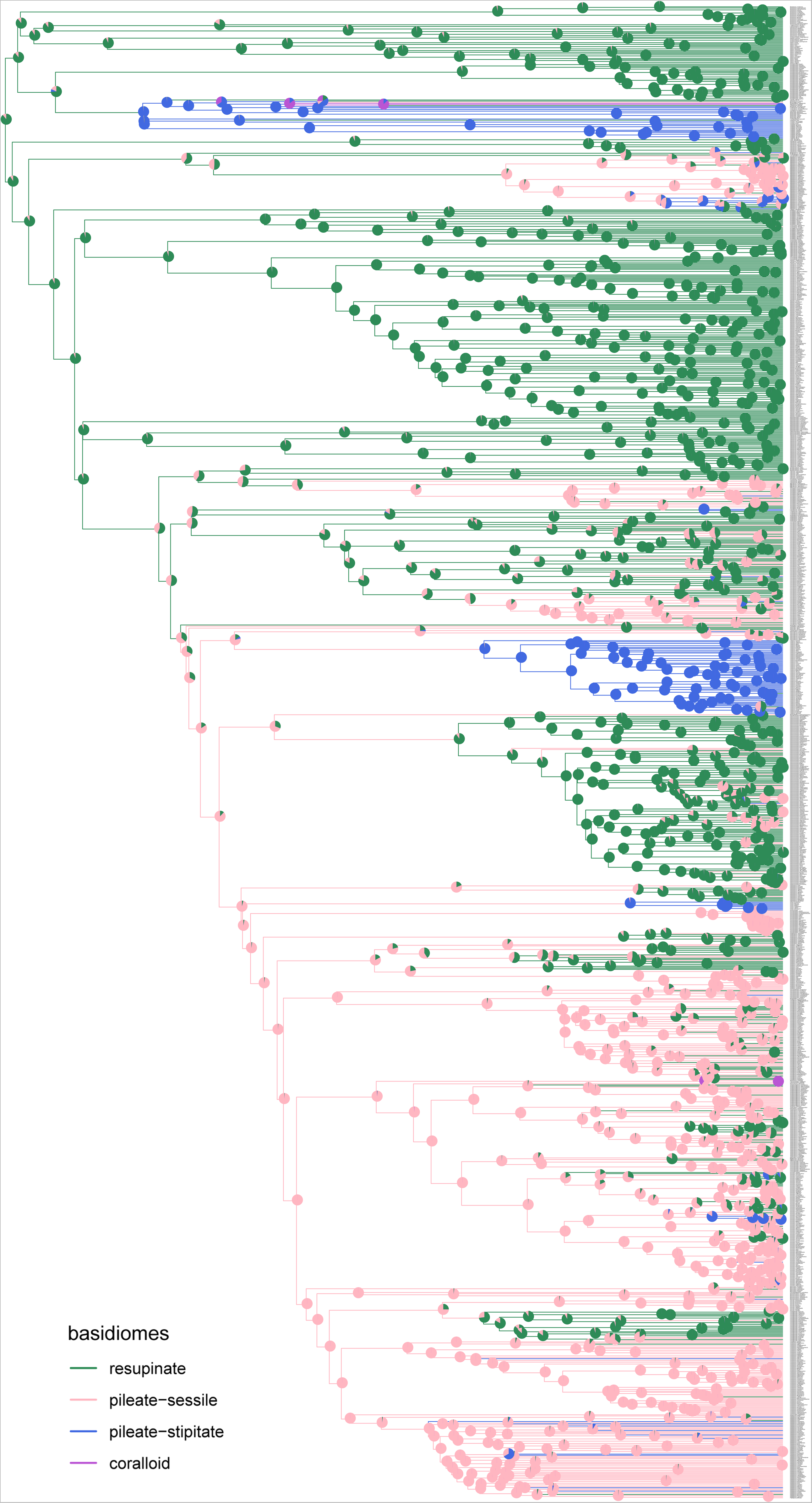
The ultrametric phylogenetic tree of *Hymenochaetales* with trait evolution of basidiomes. The posterior probability distribution for the most recent common ancestor of each node was shown as a pie chart.

**Fig. S4.**
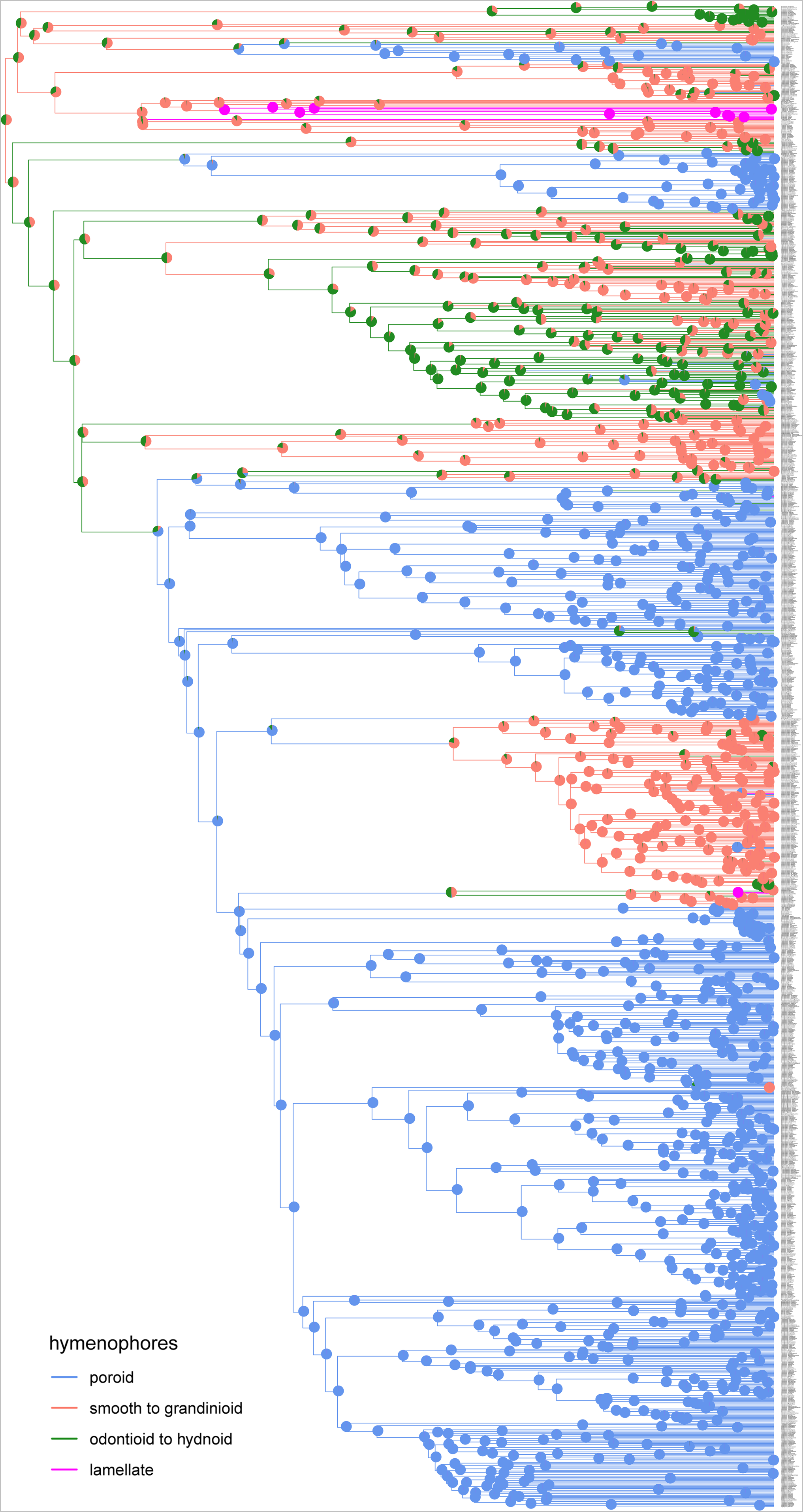
The ultrametric phylogenetic tree of *Hymenochaetales* with trait evolution of hymenophores. The posterior probability distribution for the most recent common ancestor of each node was shown as a pie chart.

**Fig. S5.**
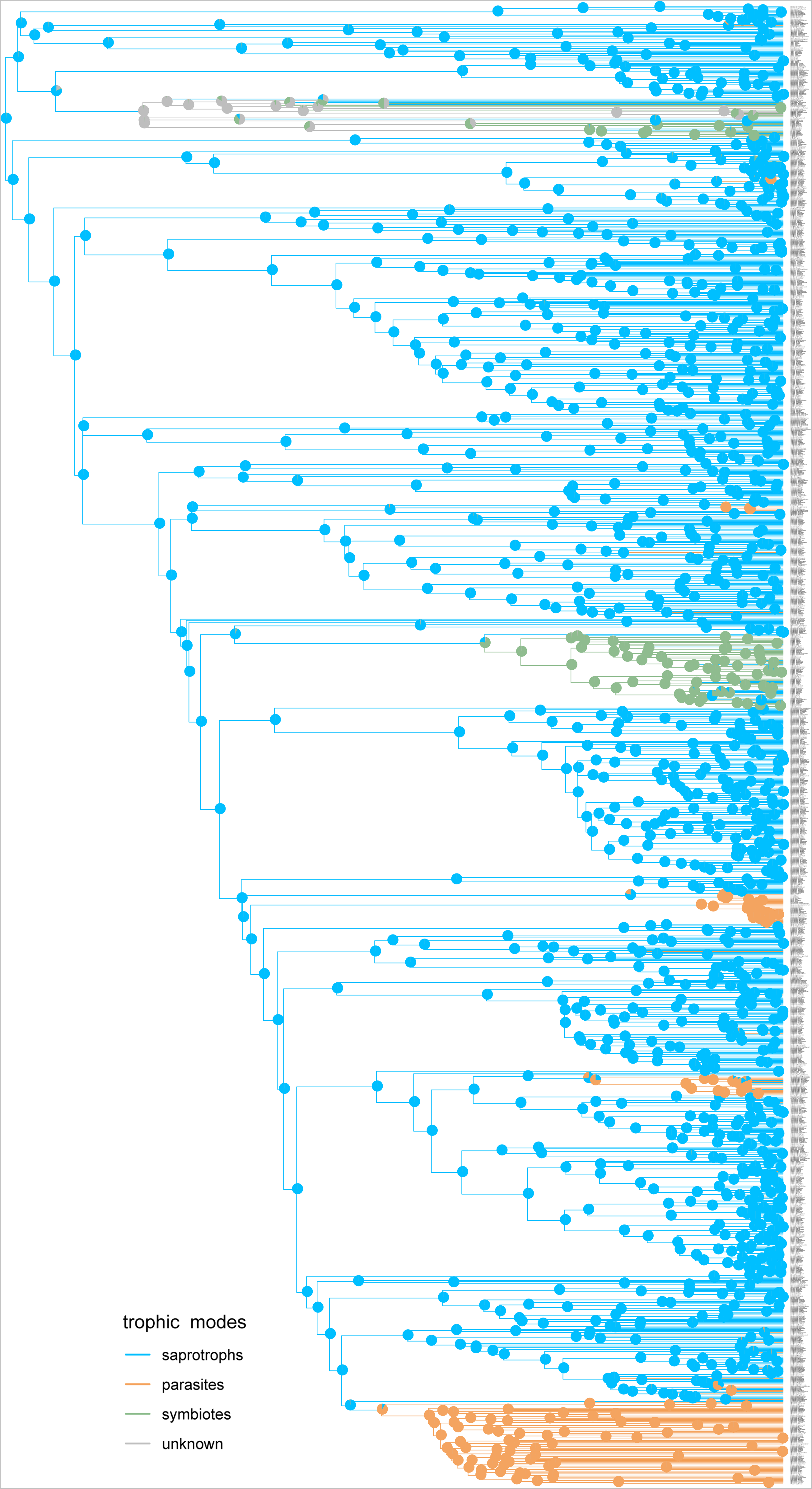
The ultrametric phylogenetic tree of *Hymenochaetales* with trait evolution of trophic modes. The posterior probability distribution for the most recent common ancestor of each node was shown as a pie chart.

**Fig. S6.**
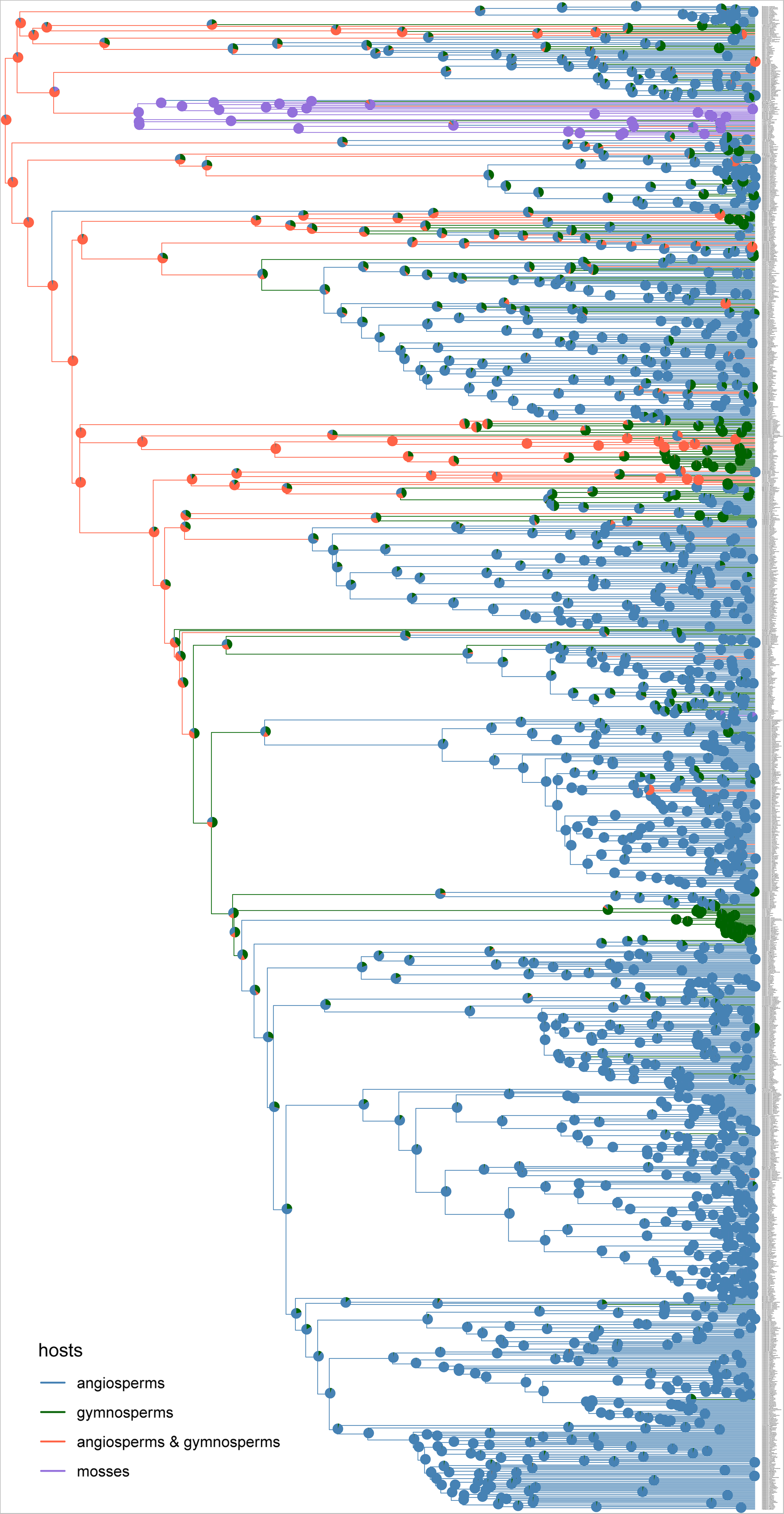
The ultrametric phylogenetic tree of *Hymenochaetales* with trait evolution of hosts. The posterior probability distribution for the most recent common ancestor of each node was shown as a pie chart.

**Fig. S7.**
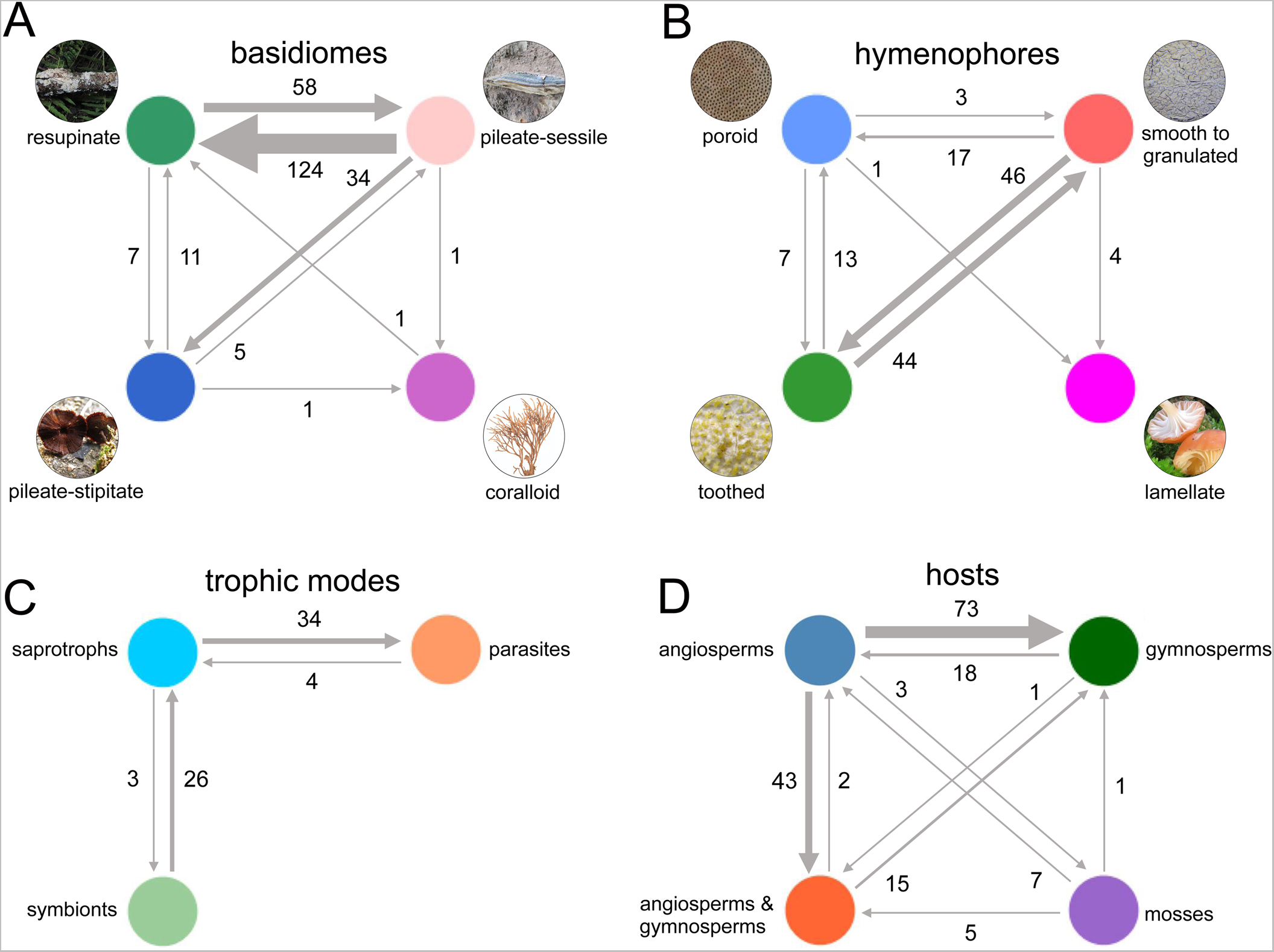
The numbers of transition events of all nodes and tips between different forms of each trait (lower left) were shown by arrows with different thicknesses; the absence of arrows between forms indicates no transition event. (**A**) basidiomes. (**B**) hymenophores. (**C**) trophic modes. (**D**) hosts.

**Fig. S8.**
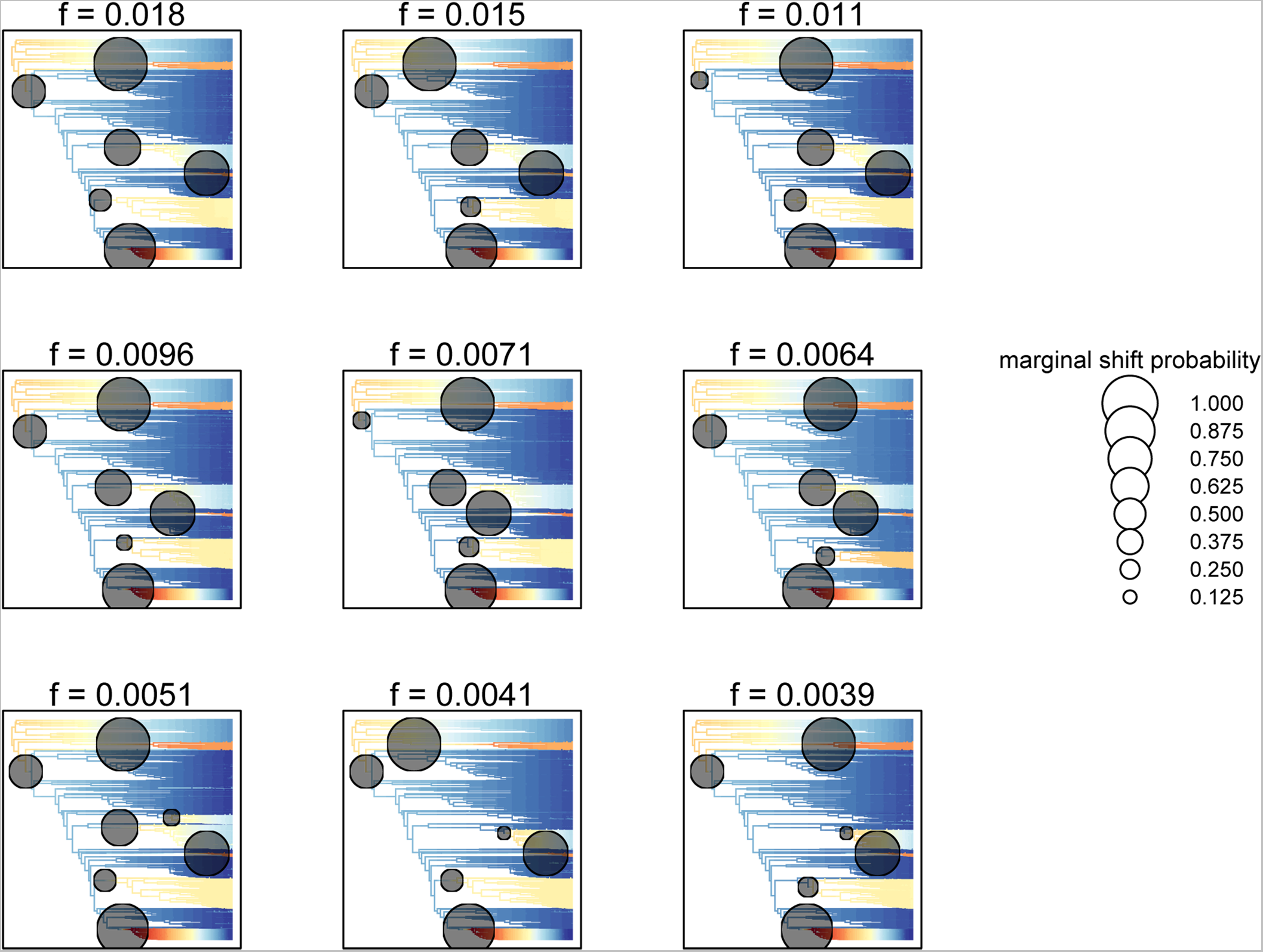
The nine most probable transition configurations inferred by Bayesian Analysis of Macroevolutionary Mixtures. These nine most probable configurations are remarkably stable with little different marginal shift probabilities.

**Fig. S9.**
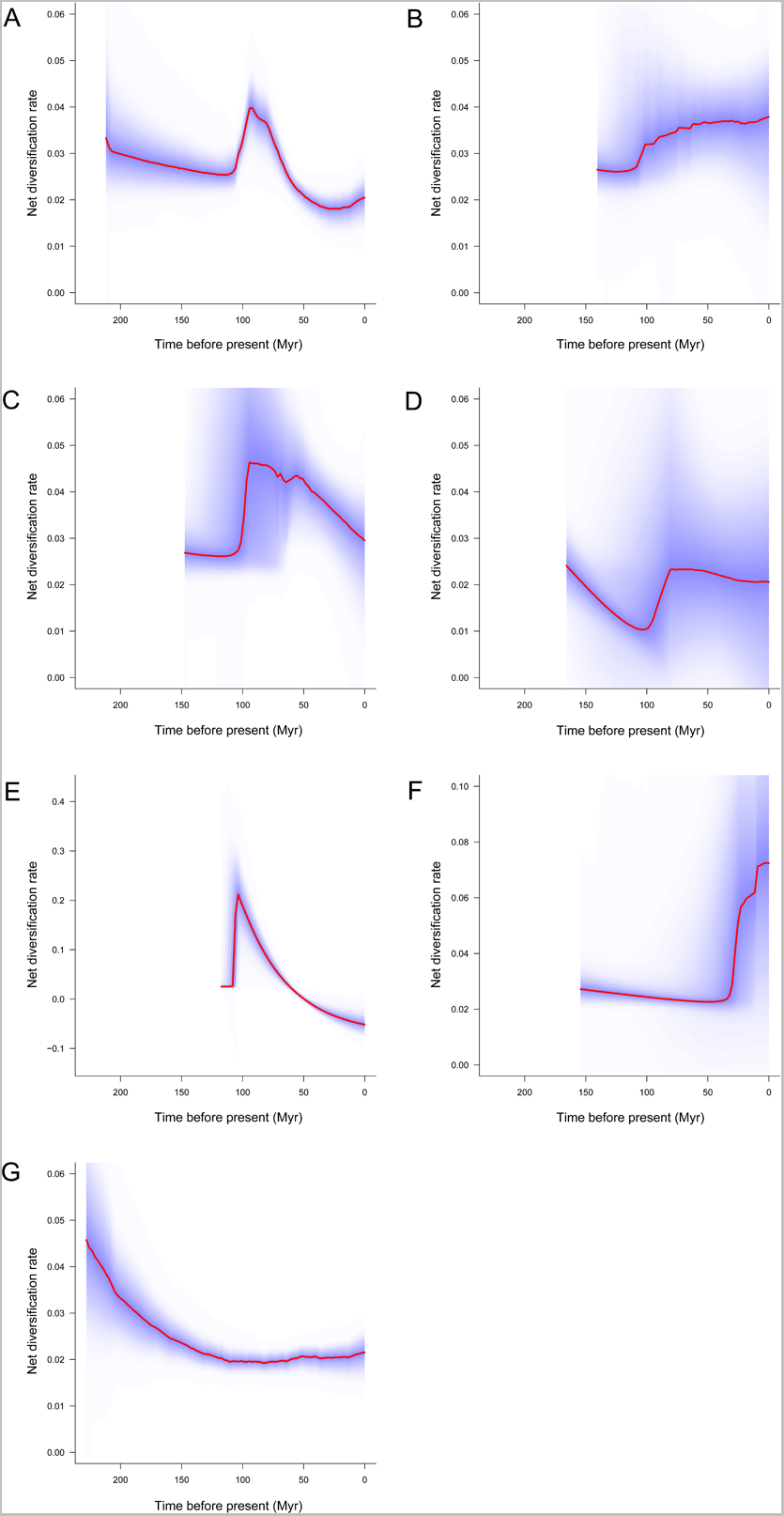
Net diversification rate through time of the six clades showing significant diversification rate shifts (**A–F**, respectively, correspond to the clades 1–6 in Fig. 3A) and the net diversification rate of *Hymenochaetales* through time after removing the clade 5 corresponding to the *Phylloporia* clade in Fig. 3A (**G**).

**Fig. S10.**
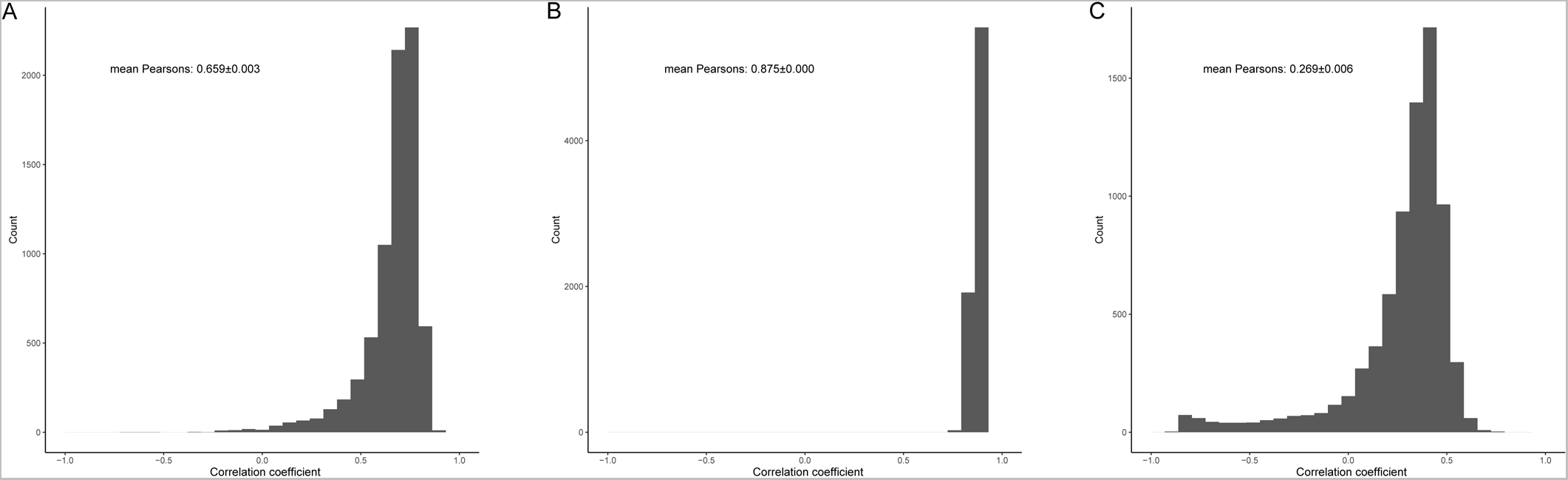
Correlation coefficients of net diversification rate and paleotemperature. (**A**) The entire *Hymenochaetales*. (**B**) The clade 5 corresponding to the *Phylloporia* clade in Fig. 3A. (**C**) The entire *Hymenochaetales* after removing the *Phylloporia* clade. The mean correlation coefficient and an estimate of the standard error inferred with Pearson’s cross-correlation analysis are shown.

**File S1** The concatenated alignment for constructing the ML phylogenetic tree.

**Table S1** Information for species used in phylogenetic and evolutionary analyses of *Hymenochaetales*.

